# Epistasis between Na^+^/K^+^-ATPase Substitutions May Influence Salinity Tolerance in *Steinernema* Entomopathogenic Nematodes

**DOI:** 10.1101/2025.07.17.665022

**Authors:** Perla Achi, Cullen McCarthy, Lanie Bavier, Robert Pena, Victoria Iglesias, Preston Christensen, Ahmed Aljidui, Anil Baniya, Connor Goldy, Rose C. Adrianza, Suraj Reddy, Simon C. Groen, Adler R. Dillman

**Author notes:** These authors contributed equally (Simon C. Groen, Adler R. Dillman). Lead contacts.

## Abstract

Soil salinity varies widely across geographies both due to natural factors and human activities, including agriculture, road salt application, sea level rise, and desertification. Increases in soil salinity may affect organisms widely and particularly impact soil foodwebs. As parasites, entomopathogenic nematodes (EPNs) occupy crucial links in soil foodwebs and are important for agriculture as biological control agents of insect pests. Previous research found that the EPN *Steinernema carpocapsae* may exhibit higher salt tolerance than several of its congeners. We recently identified that *S. carpocapsae* uniquely evolved two amino acid substitutions in the first extracellular loop of the sodium pump (Na□/K□-ATPase). Here, we tested if these substitutions explain *S. carpocapsae*’s reported lower sensitivity to salt. Our results confirm that *S. carpocapsae* exhibits higher salt tolerance and show it can more effectively locate and infect insect hosts than its congeners *S. feltiae* and *S. hermaphroditum* in highly saline environments. We then retraced the evolution of the two amino acid substitutions in *S. carpocapsae* by introducing them alone and in combination in *Caenorhabditis elegans* using CRISPR genome engineering. We found that *C. elegans* mutants with single substitutions showed improved salt tolerance. However, this improvement disappeared in the double mutant, whose sodium pump mimicked that of *S. carpocapsae*. This pattern of negative epistasis between the amino acid substitutions suggests they are not responsible for variation in salt tolerance between *Steinernema* species. Sodium pump evolution in *S. carpocapsae* might instead be driven by encounters with cardiac glycosides, which are released into soil by several clades of plants including milkweeds, sequestered by some of this EPN’s herbivorous insect hosts, and known to target the first extracellular loop of the sodium pump. Our findings provide valuable insights into EPN adaptation to changes in environmental sodium levels and may have implications for their use in biological control.

## 1. Introduction

Sodium is vital for animal development; however, its levels fluctuate widely within and across ecosystems and its availability varies for organisms occupying different trophic levels in foodwebs. The variable availability of salt influences foraging behavior and mechanisms of cellular ion transport in both vertebrate and invertebrate animals so that they may achieve homeostasis for appropriate sodium levels across tissues (Snell-Rood et al., 2014). For example, herbivorous insects often visit the soil to satisfy their physiological needs for salt (e.g., puddling in butterflies), as their plant-based diets generally contain low sodium concentrations (Xiao et al., 2010; Kaspari, 2020).

Soil salinity varies significantly depending on environmental conditions, ranging from 0 mM to 300 mM (Department of Primary Industries and Regional Development, Western Australia). Soils are classified as non-saline (<20 mM), very slightly saline (20–40 mM), slightly saline (40–80 mM), moderately saline (80–160 mM), and strongly saline (>160 mM) (Shahid et al., 2018; Stavi et al., 2021). Soil Na^+^ levels have been increasing throughout large areas of land around the globe in recent decades due to rising sea levels, climate change-related desertification, and human activities such as agriculture and road salt application (Hassani et al., 2020), posing risks to environmental sustainability and agricultural productivity (Stavi et al., 2021). This increase in salinity may pressure species to acclimate or adapt to their changing environments and may alter species interactions across multiple trophic levels in foodwebs (Nielsen et al., 2011; Snell-Rood et al., 2014).

Entomopathogenic nematodes (EPNs) fulfill important roles in plant-based multi-trophic interactions by infecting a wide range of plant-feeding (herbivorous) insects. They are found in both natural and human-modified ecosystems, where they may shape biological diversity and control insect pests. Larval and adult stages of insects become vulnerable to EPN attack while visiting the soil, which even aboveground herbivores do regularly (Achi et al., 2025a). One of the main clades of EPNs is Steinernematidae, which contains many generalist species, such as *Steinernema carpocapsae*, that can infect a variety of different insect species (Lewis and Clarke, 2012). EPNs are obligate parasites that kill their insect hosts within 48 hours of infection, playing a crucial role in controlling insect pests (Dillman et al., 2012a). Infective juveniles (IJs) initiate infection by releasing symbiotic bacteria that help overcome the host’s immune system and provide nutrients for nematode growth (Dillman et al., 2012a; Dillman and Sternberg, 2012; Lewis and Clarke, 2012; Nielsen et al., 2011; Ogier et al., 2020).

The presence of EPNs in environments is influenced by their feeding preferences, which are linked to the availability of specific host insect species (Matuska-Lyzwa et al., 2024). All EPNs engage in host-seeking behavior, though these can vary between species (Griffin, 2012; Lortkipanidze et al., 2016). Host seeking is performed by infective juveniles (IJs), which reside primarily in the soil—the IJ stage is the only one found outside of the host. IJs rely heavily on environmental cues to locate and/or remain in regions where infection is most likely; cues not only include chemicals released by plants or insect hosts such as CO□ or specialized metabolites but also temperature and vibrations (Achi et al., 2025a; Dillman et al., 2012b; Nielsen et al., 2011; Rasmann et al., 2005; Zhang et al., 2021).

Since IJs spend prolonged periods of time in the soil, they are the most likely EPNs to encounter varying salinity levels (Nielsen et al., 2011; Stuart et al., 2015). Although research on optimal environmental salt concentrations for EPNs is limited and such concentrations vary considerably among species, studies suggest that some EPNs thrive at low to moderate, but not high, salinity levels (Nielsen et al., 2011; Stuart et al., 2015). For example, increased salinity has been shown to negatively affect the foraging behavior of species such as *S. riobrave*, a generalist that deploys a ‘cruiser’ foraging strategy (Nielsen et al., 2011). Behavioral and physiological differences among EPN species might influence their responses to salt stress and may have important adaptative consequences. Such differences further raise concerns about the efficacy of EPNs as biological control agents across varying agricultural environments.

Recent studies show that *S. carpocapsae* exhibits higher salt tolerance compared to *S. feltiae* (Edmunds et al., 2021). Interestingly, *S. carpocapsae* has evolved unique substitutions (K120M, and N122H) in the first extracellular loop of the Na□/K□-ATPase (NKA) *α* subunit relative to its congeners (Achi et al., 2025a; Groen and Whiteman, 2021). The NKA is crucial for regulating ion flow, cell signaling, and stress responses (Dobler et al., 2012). Substitutions in the first extracellular loop and epistatic interactions between them have previously been linked to evolution of animal resistance to toxic cardiac glycosides (CGs; Karageorgi et al., 2019). These secondary metabolites are produced by certain plants, such as milkweeds, and sequestered by several herbivorous insects that specialize on these plants, including the monarch butterfly (Dobler et al., 2012). However, we also identified that substitutions in the first extracellular loop of the NKA found in certain herbivorous and hematophagous insects can be important for salt tolerance, which might be linked to the fact that these insects encounter relatively high sodium levels through blood feeding, feeding on salt-accumulating plants, and/or breeding in coastal or arid saline environments (Achi et al., 2025b). The presence of the substitutions in the first extracellular loop of the NKA in *S. carpocapsae* thus suggests a potential mechanism for its higher tolerance to salt stress relative to other *Steinernema* spp. In addition, other studies have suggested that stressful conditions, such as osmotic stress, may amplify the phenotypic impact of certain mutations (Katju et al., 2018).

To investigate whether the substitutions in the NKA of *S. carpocapsae* and possible genetic interactions (epistasis) between them have the potential to explain this species’ relatively high salt tolerance, we used *Caenorhabditis elegans* as a model system. The sequence of the first extracellular loop of the NKA *α* subunit is conserved across *Steinernema* spp. and *C. elegans*. Using CRISPR/Cas9 gene editing, we engineered knock-in mutant strains of *C. elegans* with the substitutions found in *S. carpocapsae* but not in its congeners. We obtained strains with these substitutions, K120M and N122H, alone and in combination, which allowed us to test for a potential role of epistasis in the evolution and maintenance of the substitutions in *S. carpocapsae*. Specifically, our study had the following aims: (1) to which extent *S. carpocapsae* displays more efficient foraging in highly saline environments relative to its congeners *S. feltiae* and *S. hermaphroditum*; (2) if *S. carpocapsae* accrues higher relative fitness than these congeners in elevated-salt conditions; (3) whether substitutions K120M and N122H that evolved uniquely in *S. carpocapsae* can contribute to salt tolerance in nematodes alone and in combination, using *C. elegans* as a model organism; and (4) if the evolution of K120M and N122H in *S. carpocapsae* has the potential to explain differences in salt tolerance among *Steinernema* spp. Understanding EPN responses to environmental cues such as sodium levels is critical, as these nematodes are widely used for biological control to target insect pests in agriculture. It could also provide deeper insights into ecological and evolutionary dynamics of multi-trophic interactions as well as molecular genetic and physiological mechanisms that enhance EPN survival and reproduction.

## 2. Materials and Methods

### 2.1. Entomopathogenic nematode strains

We cultivated nematodes of the following species in the genus *Steinernerma*: *S. carpocapsae* (All strain), *S. feltiae* (SN strain), and *S. hermaphroditum* (CS34 strain).

### 2.2. Entomopathogenic nematode white traps

Three or four waxworms (*Galleria mellonella*; Crittergrub) were placed onto p-8 filter paper in a 60-mm Petri dish. Then, 700-1,000 IJs of an EPN species were transferred into the plate. Nematodes were given four days to infect waxworms before being moved onto white traps. For this, 100-mm Petri dishes were used, each with a 30-mm Petri dish lid placed in the middle that was covered with p-5 filter paper. Waxworms were then moved onto the center of the paper and filtered tap water was used to fill enough of the larger Petri dish to get the filter paper wet along its sides but not high enough to where the water was touching the waxworms. IJs were given five to nine days to exit waxworms. Water was then collected to obtain healthy IJs and IJ suspensions were stored in cell culture flasks horizontally in a fridge at 16.7 °C.

### 2.3. Preference chemotaxis assay on agar

Chemotaxis plates were prepared following the protocol by Baiocchi et al. (2017) and left at room temperature for 12 hours before the experiment. IJ-stage EPNs were placed onto white traps to ensure we obtained fresh batches of healthy individual nematodes. The newborn IJs were then collected, washed, and prepared at a density of 4/µL. A 500-µL suspension of IJs was placed onto a waxworm for 20 minutes to stimulate contact with the host cuticle, which promotes IJ activity (Baiocchi et al., 2019). After incubation, IJs were collected, adjusted to a density of 5/µL, and prepared for chemotaxis assays. Test chemicals were prepared as follows: 2 M prenol by dissolving 203 µL of 99.9% pure prenol in 797 µL of DI water; 200 mM tetrahydrofuran (THF) by mixing 14.4 µL of THF with 985.6 µL of DI water; and 0.5 M sodium azide (NaN□) by diluting 500 µL of a 1 M stock with 500 µL of DI water. NaCl test solutions at 50-, 150-, and 300-mM concentrations were also prepared using DI water. Chemotaxis templates were printed and placed under each plate (Supplementary Fig. 1A). On one side (test #1), 5 µL of the first salt concentration was placed in the test circle, and on the opposite side (test #2), 5 µL of the second salt concentration was placed. Control plates consisted of THF (positive control for attraction) vs. DI water or prenol (negative control for repulsion) vs. DI water to verify the accuracy of the assay and that nematodes behaved as expected. THF is a known attractant for *S. carpocapsae* (Dillman et al., 2012b) and prenol a known repellant for both *S. carpocapsae* and *S. feltiae* (Baiocchi et al., 2017; Baiocchi et al., 2020). Test plates compared NaCl concentrations of 50 vs. 150 mM, 50 vs. 300 mM, and 150 vs. 300 mM. Chemicals were allowed to settle for 15 mins before adding 2 µL of 0.5 M NaN□ to each scoring circle. A 15-µL suspension of IJs at a density of 5/µL (70–170 nematodes total) was then placed on opposite sides in a diamond pattern, alternating between test solutions and nematodes. Plates were stacked in random groups of three with alternating orientations and placed in a box with a lid on a vibration-resistant platform. The assay ran for 24 hours, after which the location of the nematodes—test #1, test #2 or neither—was recorded. Choice percentages were calculated by counting the number of nematodes in each test area, dividing by the total number of IJs inoculated, and multiplying by 100. The experiment was performed in triplicate within each of three biological replicates. Data were graphed using means across biological replicates, and statistical analysis was conducted using two-way ANOVA in PRISM, with results presented as mean ± SEM.

### 2.4. Preference chemotaxis assay in sand

Sand was autoclaved and then repeatedly washed with tap water followed by DI water. The sand was dried overnight at 60 °C before use. An olfactometer was set up according to the design in Supplementary Fig. 1B and 1C (Wu and Duncan, 2020) and measurements of the pipes and glass tubes were recorded. Sand was moistened to 12% with either tap water (control) or NaCl test solutions at concentrations of 0, 50, 150, and 300 mM. A consistent weight of sand (28 g) per tube was used for each replicate and trial. To eliminate bias, the central region of the olfactometer was filled with well-packed control sand moistened to 12% to create a neutral zone. A total of 1,000 IJs in 100 µL of water were released into the center of the olfactometer, ensuring they were placed beneath the soil surface to prevent them from experiencing a dry environment. IJs were collected from fresh white traps and stimulated by exposing them to waxworms prior to the experiment as described above. The olfactometer was placed horizontally on foam to prevent vibration, kept in the dark, and randomly oriented to avoid directional bias. The assay ran for 24 hours, after which the tubes from each side were removed separately. Nematodes were then collected using the Baermann funnel technique and their densities were measured. The location of the nematodes was recorded as either the test area, the control area, or the middle. Choice percentages were calculated by counting the number of nematodes in the control or test area, dividing by the total number of IJs inoculated, and multiplying by 100. The experiment was conducted in triplicate within each of three biological replicates. Data were graphed in PRISM using means across biological replicates, and chi-squared analysis was performed on the average of each replicate, with the number of nematodes in the control area as the expected value and the number of nematodes in the test area as the observed value.

### 2.5. Infectivity and foraging behavior assay in sand

EPNs were tested on their ability to find and infect a host successfully under conditions differing in salinity. Containers consisting of moistened sand (12%) with either 0, 50, 150 or 300 mM NaCl were prepared. Three healthy waxworms were buried in the sand on one side of the container using cheese cloth (Supplementary Fig. 1D). One thousand IJs in 100 µL of water were released onto the opposite side of the container, about 10 cm away from the waxworms. Care was taken to place the nematodes in the soil and not on the surface to prevent them from experiencing dehydration. The waxworms were collected 48 hours later and dissected. EPN infectivity was measured by counting the number of EPNs found inside each waxworm. EPN foraging ability was measured by assessing indicators of infection such as EPN presence in waxworms. Foraging was graphed by the proportion of waxworms found to be infected. Measurements of the container (13 cm x 8 cm x 6.5 cm) were taken and a consistent weight of sand (260 g) was used in each replicate. The experiment was performed five times to obtain a sufficient sample size. Data were graphed using means across replicates, and statistical analysis was conducted using two-way ANOVA in PRISM, with results presented as mean ± SEM.

### 2.6. Media preparation and bacterial maintenance

#### Thick nematode growth media (NGM)

Thick NGM was prepared using 3 g of NaCl, 20 g of agar, 2.5 g of peptone, 2 g of dextrose, 975 mL of DI water, and 10 mL of uracil. After autoclaving, 25 mL of sterile potassium phosphate buffer, 1 mL of filtered magnesium sulfate, and 1 mL of filtered calcium chloride were added under sterile conditions. Subsequently, 1 mL of cholesterol was added. The media was poured into 100-mm Petri dishes, left to dry at room temperature for 48 hours, and then stored in a refrigerator at 4 °C.

#### LB+SP agar

LB+SP agar was made with 40 g of LB agar, 1 g of sodium pyruvate, and 1 L of DI water, then autoclaved.

#### LB + SP broth

LB+SP broth was made with 25 g of LB broth, 1 g of sodium pyruvate, and 1 L of DI water, then autoclaved.

#### Bacterial maintenance

*Escherichia coli*, strain OP50, was used to maintain *C. elegans* nematodes. OP50 bacteria were streaked weekly on LB+SP agar in 100-mm Petri dishes, left overnight in an incubator at 28 °C, and stored in a 20-°C incubator. Single colonies were transferred into a 15-mL conical tube that contained ∼8 mL of LB+SP broth. Bacteria were left to incubate in a shaker at 28 °C overnight, then stored in a 20-°C incubator.

### 2.7. CRISPR editing of *ATPα* in *Caenorhabditis elegans*

Gene editing of *ATPα* (*eat-6*), the gene encoding the NKA*α* subunit, in *C. elegans* was performed by Suny Biotech using CRISPR-Cas9 technology. Four gene-edited strains were engineered in the N2 background: a CRISPR control strain with three synonymous mutations in *eat-6* [eat-6(t502g,t508c,c520t)-N2], two strains with additional single non-synonymous mutations in *eat-6* that led to single amino acid substitutions in the NKA*α* subunit [eat-6(K105M)-N2 and eat-6(N107H)-N2] and a strain with both additional non-synonymous mutations in *eat-6* combined [eat-6(K105M, N107H)-N2]. The amino acid substitutions are referred to here using the protein residue numbering for the mature *C. elegans* NKA*α* subunit but are referred to throughout the rest of this study using the protein residue numbering for the mature pig (*Sus scrofa*) protein to facilitate comparisons with previous phylogenetic studies of ATPα in insects and vertebrates. Accordingly, the *C. elegans* strains are named here based on their amino acid residues at positions orthologous to positions 111, 119, 120, and 122 of the mature pig NKA*α* subunit (positions 96, 104, 105, and 107 in *C. elegans*, respectively). The wild-type *C. elegans* strain N2 is referred to as DSKN, the CRISPR-control strain as DSKN^C^, strain eat-6(K105M)-N2 as DSMN, strain eat-6(N107H)-N2 as DSKH, and strain eat-6(K105M, N107H)-N2 as DSMH (Table 1; Supplementary Table 1). Positions 111 and 119 are included in our naming convention since amino acid substitutions at these positions have previously been implicated in salt tolerance differences among insects (Achi et al., 2025b).

**Table 1.**
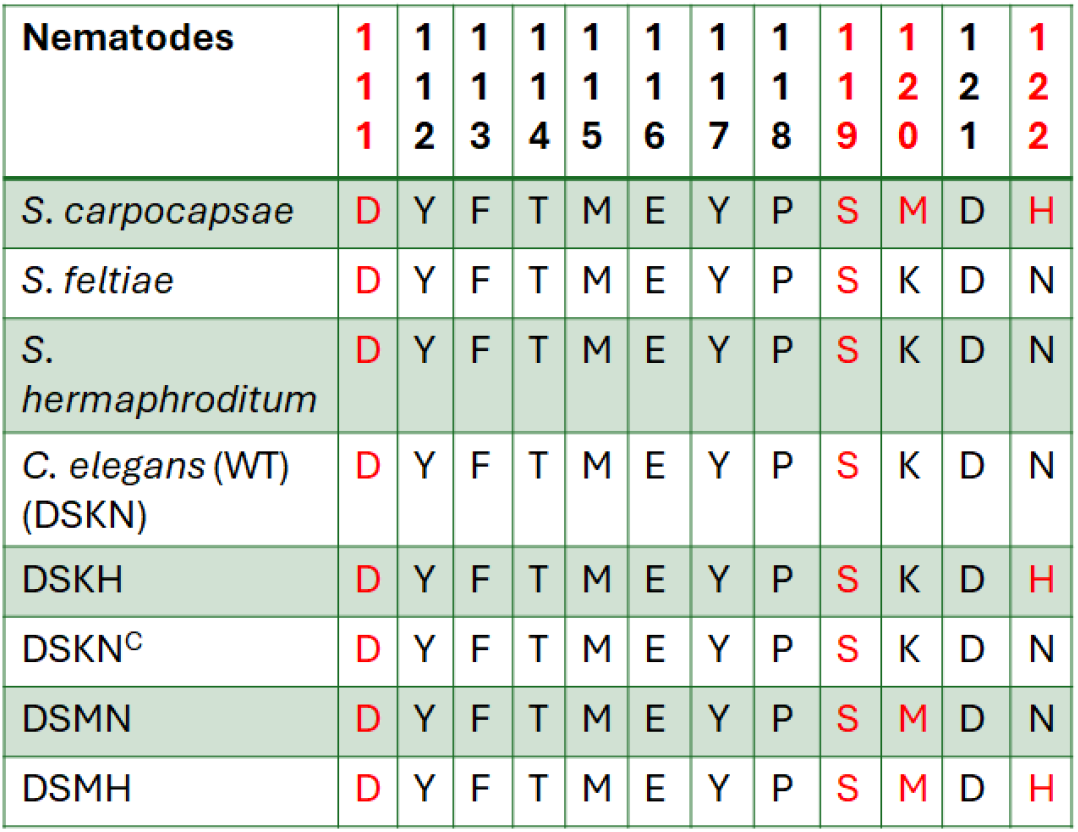
Amino acid sequences of the Na□/K□-ATPases of *Steinernema* spp. and *Caenorhabditis elegans* (N2 wild-type [WT / DSKN]) and mutant strains ([DSKN^C^], [DSKH], [DSMN], and [DSMH]). The sequences at positions 111-122 of the Na□/K□-ATPase α subunit are shown (numbering relative to the pig ATP1A1 sequence). The names of the *C. elegans* strains refer to their amino acid residues at positions 111, 119, 120, and 122 with DSKN^C^ referring to the CRISPR-control strain.

### 2.8. Maintenance and culturing of *Caenorhabditis elegans*

OP50 bacteria were grown in a 15-mL conical tube with LB+SP broth. Thick NGM plates were spotted with bacteria from this broth a day before *C. elegans* nematodes were added for culturing. Nematodes were moved to fresh media with OP50 bacteria weekly to ensure optimal growth conditions. Once the nematodes were transferred onto new plates, the plates were left at room temperature.

### 2.9. Synchronization of *Caenorhabditis elegans*

Adult stages of *C. elegans* were collected from plates using M9 buffer and placed into a 15-mL conical tube. Nematodes were centrifuged at 1 rpm for 1 min. Multiple collections from plates were performed until a nematode pellet of 0.5-1 mL was obtained. Bleaching solution was prepared by adding 0.16 g of NaOH, 2 mL of 5% bleach, and 8 mL of DI water to a 15-mL conical tube. From this, 6-7 mL were then added to the conical tube with the nematode pellet. The conical tube was shaken vigorously for three mins. M9 buffer was added immediately afterwards to stop the reaction. The conical tube was then centrifuged and washed three times with M9 buffer, leaving only the nematode pellet at the bottom and ensuring removal of all bleaching solution. Nematodes were then left in 3 mL of M9 buffer in the conical tube on a rotator overnight to allow egg hatching by the next day.

### 2.10. Salt tolerance assay with entomopathogenic nematodes and *Caenorhabditis elegans*

Nematodes were placed in solutions of NaCl and KCl to investigate their ability to tolerate varying environmental conditions (Khanna et al., 1997; Supplementary Fig. 3A). Three solutions were used in this experiment: K-medium as the control solution, along with NaCl and KCl test solutions. K-medium was prepared in a 1-L Erlenmeyer flask with 51.3 mM (3.0 g) NaCl, 31.7 mM (2.36 g) KCl, and 1,000 mL of DI water. Using the same ingredients, but adjusting the concentrations of either NaCl or KCl, the saline test solutions were prepared. The NaCl solutions were made at concentrations of 265.2 mM (15.5 g) and 282.3 mM (16.5 g), while the KCl solutions were prepared at 155.1 mM (11.6 g) and 169.5 mM (12.6 g). All three solutions were autoclaved and next, 1 mL of each control or test solution was added to individual wells of 12-well plates that contained a thin layer of thick NGM. Approximately 5-25 adult *C. elegans* nematodes were then placed in each well. For EPN assays, IJs were stimulated with waxworms for 20-25 minutes before transferring 5-25 of them per well. Plates were incubated at 20 °C for 24 hours, after which lethality was assessed by gently poking worms with a titanium brush to confirm immobility. For *C. elegans*, a percent lethality below 10% was considered normal (Khanna et al., 1997). Salt tolerance assays were conducted only at 265.2 mM NaCl and 155.1 mM KCl because *C. elegans* showed sensitivity to higher concentrations. All experiments were performed in triplicate. Graphs were generated using PRISM to display the mean ± SEM, and statistical analysis was performed using two-way ANOVA.

### 2.11. Salt fitness assay with entomopathogenic nematodes and *Caenorhabditis elegans*

To investigate nematode fecundity and survival across different salt levels, thick NGM plates were prepared, including control plates with 60 mM NaCl and highly saline plates with 200, 250, and 300 mM NaCl (Supplementary Fig. 4A). The saline NGM plates were made using the same ingredients as the control plates but with adjusted NaCl measurements to achieve the final concentrations. The day before the experiment, each well in a set of 24-well plates was spotted with 15 µL of OP50 bacterial suspension. On the day of the experiment, a single L1 *C. elegans* larva was placed into each well. For each *C. elegans* strain (DSKN, DSKN^C^, DSMN, DSKH, and DSMH), one control plate and one saline NGM plate were used per round, resulting in 10 plates per round (5 regular and 5 saline). The experiment was conducted over three rounds. Nematode progeny production was monitored from days 3 to 6. On day 6, new plates were prepared by spotting wells with OP50 bacteria. On day 7, a single larva from a well of an old plate was transferred to a well of a new plate with the same NaCl concentration, allowing progeny production to be monitored for a second generation. Progeny counts and survival were recorded from days 3 to 6 on the second set of plates. Graphs were generated using PRISM to display the mean ± SEM, and statistical analysis was performed using two-way ANOVA for fecundity and log-rank analysis for survival.

### 2.12. Graphing and statistical analysis

All graphs were made using PRISM with the mean and standard error of the mean (SEM). Depending on data type, statistical analysis was performed using log-rank analysis, chi-squared analysis or two-way ANOVA that was followed by post hoc Tukey’s HSD tests. Figures were assembled using Adobe Photoshop and illustrations were created using BioRender.

## 3. Results

### 3.1. *Steinernema carpocapsae* preferred intermediate salt concentrations on agar

IJs of EPN species *S. carpocapsae, S. feltiae*, and *S. hermaphroditum* were tested on their preference for different NaCl concentrations using a chemotaxis behavioral assay (Supplementary Fig. 1A). THF was used as a positive control for EPN attraction and prenol as a negative control, since it is known to repel *S. carpocapsae* and *S. feltiae* (Baiocchi et al., 2017; Baiocchi et al., 2020). IJs of *S. carpocapsae* showed significant attraction to THF when tested against DI water (P=0.0038). While IJs of *S. hermaphroditum* and *S. feltiae* exhibited a trend toward attraction, this was not significant (Supplementary Fig. 2A). All three EPN species demonstrated significant repulsion to prenol (P<0.001), indicating that the EPNs were behaving normally and that the assay worked as expected (Supplementary Fig. 2B).

Among the three EPN species, *S. carpocapsae* displayed a significant preference for 150 mM NaCl when tested against 50 mM (P=0.0382), whereas *S. feltiae* showed a significant preference for 50 mM NaCl in the same assay (P=0.0298; Fig. 1A). No significant differences in EPN preference were observed across the other comparisons. However, IJs of *S. feltiae* showed a bias for migrating towards 50 mM NaCl in the 50 mM vs. 300 mM assay, and both *S. carpocapsae* and *S. feltiae* trended towards a preference for 150 mM in the 150 mM vs. 300 mM assay (Fig. 1B, 1C). IJs of *S. hermaphroditum*, on the other hand, showed no preferences or trends towards these with any of the combinations of salt concentrations tested.

**Fig. 1:**
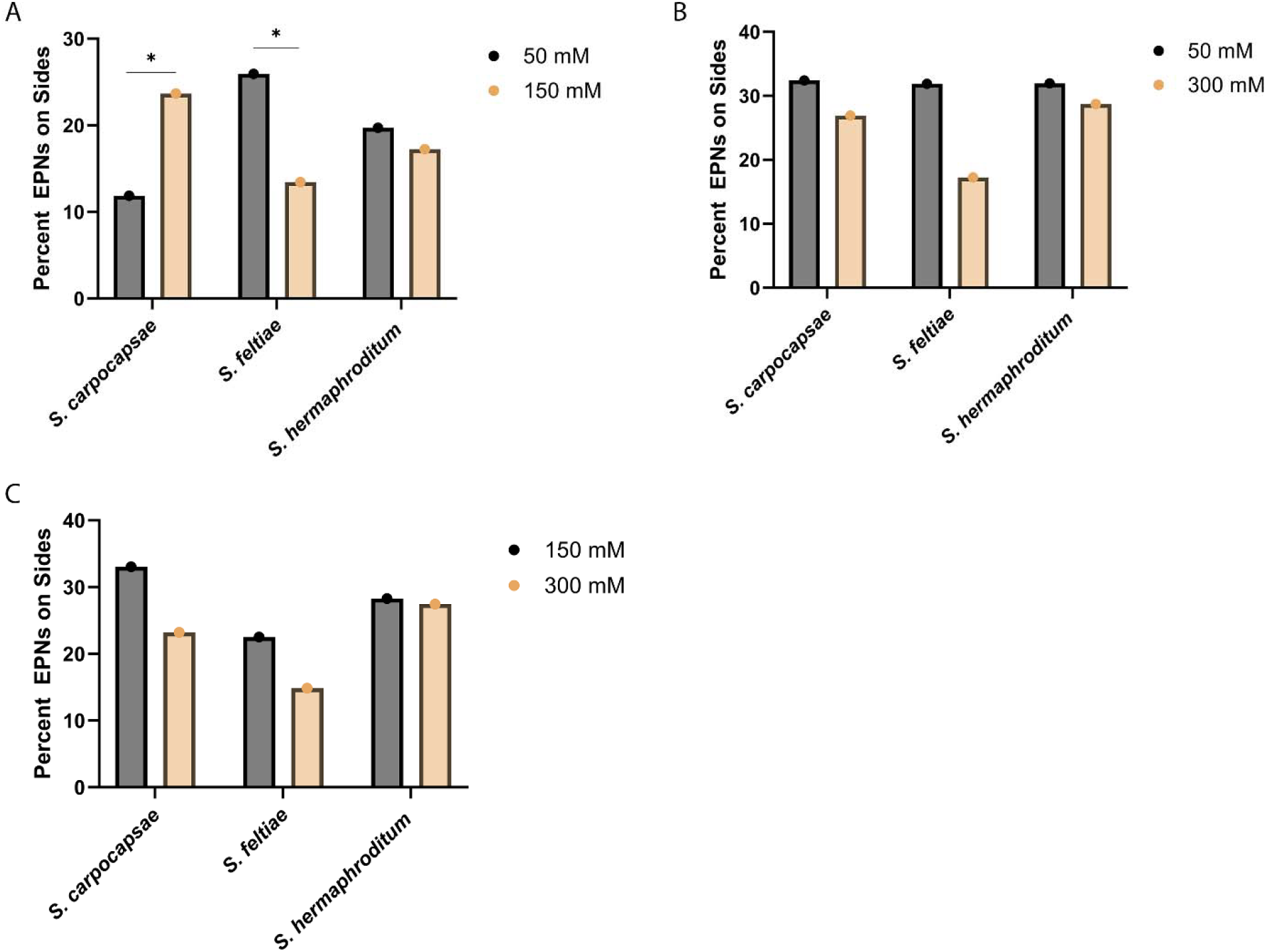
*Steinernema carpocapsae* demonstrates a preference for intermediate salt concentrations on agar plates. **A-C)** Percentage of EPN infective juveniles found on each side of a two-choice behavioral assay (50 vs. 150 mM NaCl, 50 vs. 300 mM NaCl or 150 vs. 300 mM NaCl, respectively). The X axis represents the different EPN species and the Y axis represents the relative number of nematodes (%) found on either side. Data is represented as means. Statistical analysis was performed using two-way ANOVA followed by post-hoc Tukey’s HSD tests in PRISM.

### 3.2. *Steinernema carpocapsae* preferred relatively high salt concentrations in sand

Next, EPNs were tested for movement towards areas with different salt concentrations in sand (50 mM, 150 mM, and 300 mM), following the experimental design outlined in Supplementary Fig. 1B and 1C. The nematodes were first given a control-control choice option (0 mM vs. 0 mM NaCl) to confirm they had no directional bias, with approximately 50% of the EPNs expected to choose each side (Fig. 2A-C). Once this was confirmed, EPN preferences for different salt levels were tested.

**Fig. 2:**
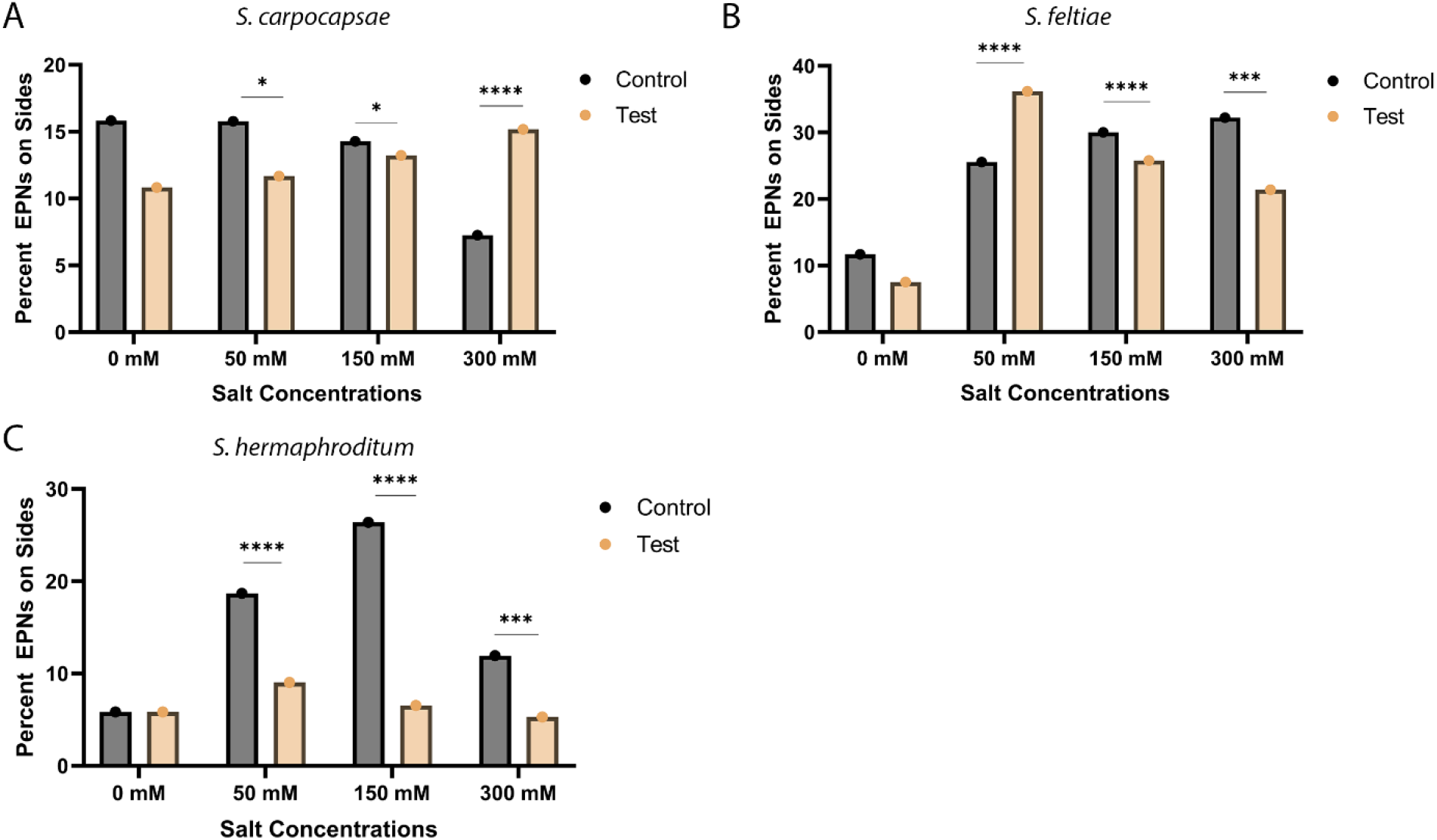
*Steinernema carpocapsae* has a significant preference for 300 mM NaCl in sand. **A-C)** Percentage of *S. carpocapsae, S. feltiae*, and *S. hermaphroditum* (respectively) infective juveniles in a two-choice behavioral assay navigating to a test side with salt concentrations of 0, 50, 150 or 300 mM vs. the control side. Data is represented as means. Statistical analysis was performed using chi-squared tests with the control side serving as the expected outcome and a test side serving as the observed outcome.

IJs of *S. carpocapsae* showed a significant preference for the control in assays comparing it to 50 mM and 150 mM NaCl (P=0.0142 and P=0.0234, respectively). However, IJs of this species exhibited a significant preference for 300 mM NaCl over the control (P<0.0001; Fig. 2A). IJs of *S. feltiae* demonstrated a significant preference for 50 mM NaCl over the control (P<0.0001) but shifted to preferring the control significantly as salt levels increased (P<0.0001; Fig. 2B). IJs of *S. hermaphroditum* preferred the control over the NaCl test concentrations in all assays (P≤0.001; Fig. 2C). Taken together, these data suggest that *S. carpocapsae* consistently prefers higher salt concentrations than *S. feltiae* and *S. hermaphroditum*.

### 3.3. *Steinernema carpocapsae* was able to locate and infect insects in highly saline conditions

Next, the EPNs were assessed for their ability to seek and infect a waxworm host at the same salt concentrations tested in the preference assays, using the setup described in Supplementary Fig. 1D. No significant differences were observed between IJs of the three EPN species in their ability to locate a host across the different salt concentrations (Fig. 3A). However, when analyzing the number of EPNs successfully infecting each waxworm, *S. hermaphroditum* exhibited significantly reduced infection success at 300 mM NaCl when compared to 50 mM (P=0.0027; Fig. 3B). Similarly, *S. feltiae* demonstrated a significant reduction in its ability to infect a host starting at 150 mM NaCl (P=0.0437), with even greater difficulties at 300 mM (P=0.0002). This suggests that while these species can still locate hosts at salt concentrations higher than 50 mM, they are less able to successfully infect them.

**Fig. 3:**
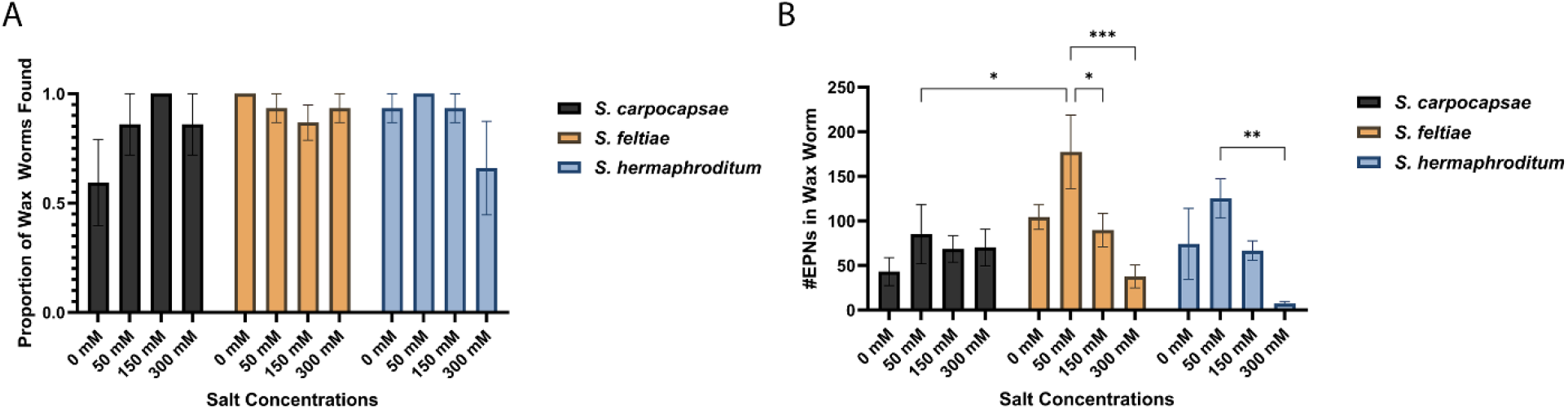
*Steinernema carpocapsae*’s infectivity of insect hosts does not decline at 300 mM NaCl. **A)** Proportion of waxworms that were located by infective juveniles of *S. carpocapsae, S. feltiae*, and *S. hermaphroditum* at salt concentrations of 0, 50, 150, and 300 mM in sand. **B)** Number of EPNs found per waxworm at each salt concentration. Data are represented as means ± SEM. Statistical analysis was performed using two-way ANOVA followed by post-hoc Tukey’s HSD tests in PRISM.

Notably, *S. carpocapsae* showed no significant differences in infection success regardless of the salt concentrations tested (Fig. 3B). For all three species, although not statistically significant, the highest infection rates were observed at 50 mM NaCl, suggesting that some salt in the environment may benefit the infective capabilities of EPNs. Interestingly, at this concentration, significantly higher numbers of EPNs infected each waxworm for *S. feltiae* than for *S. carpocapsae* (P=0.0163). Taken together, these data indicate that while *S. feltiae* and *S. hermaphroditum* continue to be highly infectious at salt concentrations of up to 150 mM, *S. carpocapsae* exhibits higher salt tolerance and retains its ability to successfully infect hosts in environments exceeding NaCl levels of 150 mM.

### 3.4. Na^+^/K^+^-ATPase substitutions K120M and N122H do not explain the higher salt tolerance of *Steinernema carpocapsae*

Amino acid substitutions in the first extracellular loop of the NKA, particularly at positions 111 and 119, have previously been shown to influence salt tolerance in insects (Achi et al., 2025b). Since *S. carpocapsae* evolved unique amino acid substitutions in the first extracellular loop of the NKA relative to other *Steinernema* spp. (K120M and N122H), we aimed to assess whether these could contribute to the higher salt tolerance observed for *S. carpocapsae*. As it is unclear in which order the substitutions evolved in the lineage leading up to *S. carpocapsae*, we further wished to find out if their evolution could have been shaped by epistasis. Using CRISPR gene editing, several *C. elegans* knock-in mutant strains were engineered, which are referred to by their amino acid residues at positions 111, 119, 120, and 122. The experimental design of the salt tolerance assay that the *C. elegans* mutant strains and *Steinernema* spp. EPNs were subjected to is outlined in Supplementary Fig. 3A. Nematodes were exposed to a control buffer (K-medium) or various concentrations of NaCl and KCl solutions, as informed by previous research (Fig. 4, Supplementary Fig. 3; Khanna et al., 1997).

**Fig. 4:**
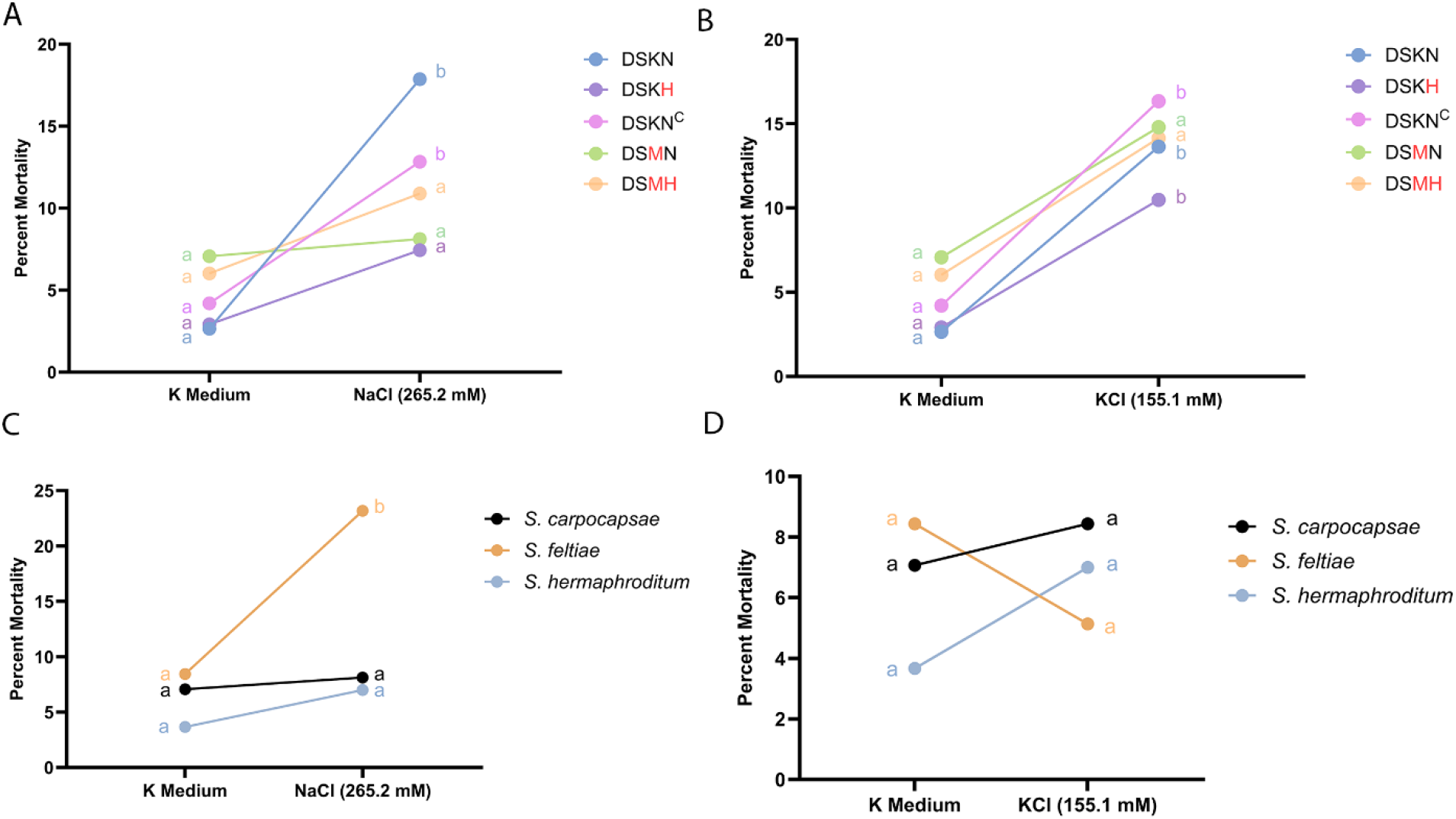
Na^+^/K^+^-ATPase substitutions K120M and N122H significantly enhance salt tolerance in nematodes. **A)** Percent mortality of *Caenorhabditis elegans* strains DSKN, DSKN^C^, DSKH, DSMN, and DSMH (described in Table 1) at 24 hours post exposure to K-medium buffer or 265.2 mM NaCl. **B)** Percent mortality of each *C. elegans* strain at 24 hours post exposure to K-medium buffer or 155.1 mM KCl. **C)** Percent mortality of *Steinernema carpocapsae, S. feltiae*, and *S. hermaphroditum* at 24 hours post exposure to K-medium buffer or 265.2 mM NaCl. **D)** Percent mortality of each *Steinernema* species at 24 hours post exposure to K-medium buffer or 155.1 mM KCl. Data is represented as means. Statistical analysis was performed using two-way ANOVA followed by post-hoc Tukey’s HSD tests in PRISM.

Initially, the DSKN (N2 wild-type), DSKN^C^ (CRISPR-control), and DSKH strains of *C. elegans* were exposed to salt solutions for 24 hours to determine the optimal concentrations for further testing. All three strains exhibited less than 10% mortality in the K-medium buffer, which is considered normal (Khanna et al., 1997). At 265.2 mM NaCl, DSKH displayed less than 10% mortality, which was similar to its mortality in the buffer solution (Fig. 4A). However, both DSKN and DSKN^C^ showed mortality rates above 10% in this solution, which was significantly higher than in the K-medium (P<0.0001 and P=0.0175, respectively). When the NaCl concentration was increased to 282.3 mM, no significant differences were found among the strains, although all three showed an increase in mortality to above 10%. Therefore, higher NaCl concentrations were not tested (Supplementary Fig. 3B). The two additional *C. elegans* knock-in mutant strains (DSMN and DSMH) were also tested at 265.2 mM NaCl. DSMN and DSMH responded similarly to this salt concentration as DSKH, demonstrating NaCl tolerance at 265.2 mM, with no significant differences in mortality relative to the K-medium (Fig. 4A). The nematodes were further exposed to 155.1 mM and 168.5 mM KCl. In both solutions, DSKN, DSKN^C^, and DSKH showed a significant increase in mortality (above 10%) compared to the K-medium (Fig. 4B, Supplementary Fig. 3C). This increase was not observed for DSMN and DSMH (Fig. 4B), although their mortality remained above 10%, indicating some sensitivity.

The assays were then repeated with the EPN species: *S. carpocapsae* (P=0.0023) and *S. hermaphroditum* (P=0.0013) exhibited significantly lower mortality compared to *S. feltiae* at 265.2 mM NaCl (Fig. 4C). Indeed, *S. feltiae* had a significantly negative response to this NaCl concentration when compared to K-medium (P=0.0010), unlike the other EPN species. At 155.1 mM KCl, no EPN species showed a significant difference in mortality when compared to the K-medium (Fig. 4D) and all three species had mortality rates below 10%, suggesting a general tolerance to these KCl levels.

Taken together, our results indicate that *S. carpocapsae* and *S. hermaphroditum* have a higher NaCl tolerance in this assay than *S. feltiae*. In addition, negative epistasis may exist between substitutions K120M and N122H that evolved uniquely in *S. carpocapsae*, since each substitution on its own improved salt tolerance in *C. elegans* but no additional improvement was observed when both substitutions were present together. When combined, these observations further suggest that, although substitutions K120M and N122H might contribute to *S. carpocapsae*’s enhanced salt tolerance, they cannot explain why both *S. carpocapsae* and *S. hermaphroditum* were less sensitive to salt in terms of survival than *S. feltiae*.

### 3.5. Na^+^/K^+^-ATPase substitutions K120M and N122H alone, but not in combination, allow for greater *Caenorhabditis elegans* fecundity in highly saline conditions

The results of the previous salt tolerance assays suggested the existence of negative epistasis between substitutions K120M and N122H that evolved along the lineage leading up to *S. carpocapsae*. To confirm patterns of epistasis between these substitutions, we monitored both survival and fecundity fitness of the *C. elegans* knock-in mutants for two generations at different salt concentrations. No significant differences in survival were observed between the strains in generation 1 (^1^) or generation 2 (^2^; Supplementary Fig. 4B, 4C). In terms of fecundity, all strains displayed similar patterns in the first generation, showing a significant decrease in fecundity between the control (60 mM NaCl) and all other salt concentrations on day 6 of the assay. The strains with the wild-type NKA sequences also showed a significant decrease in fecundity at the higher concentrations of 250 and 300 mM NaCl relative to the 200 mM solution during the first generation (P=0.0140 and P=0.0259 for DSKN^1^ at 250 and 300 mM, respectively; P=0.0454 and P=0.0289 for DSKN^C1^ at 250 and 300 mM, respectively; Fig. 5A, 5C, and 5F). DSMH^1^ showed no significant difference in fecundity between 200 and 250 mM or 250 and 300 mM but experienced a significant decrease in fecundity at 300 relative to 200 mM (P=0.0097; Fig. 5E, 5F). DSKH^1^ and DSMN^1^ showed changes in fecundity that were similar to one another, displaying a significant decrease in fecundity for the 200, 250, and 300 mM NaCl solutions relative to the control solution (60 mM) but no significant differences between the 200, 250 or 300 mM NaCl solutions (Fig. 5B, 5D, 5F).

**Fig. 5:**
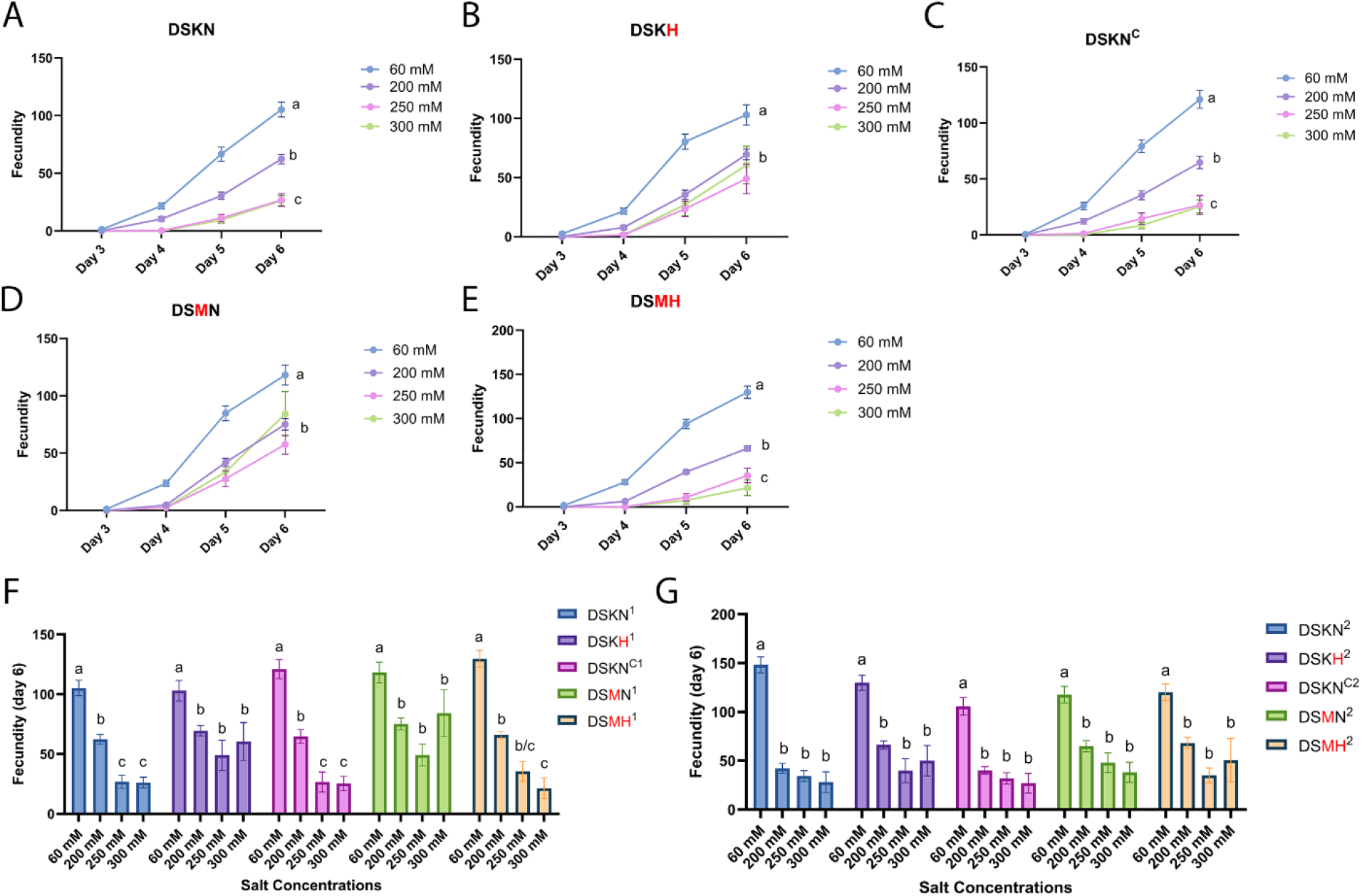
Na^+^/K^+^-ATPase substitutions K120M or N122H individually conferred the strongest tolerance to highly saline conditions when considering nematode fecundity. **A-E)** Fecundity of *Caenorhabditis elegans* strains DSKN, DSKN^C^, DSKH, DSMN, and DSMH (described in Table 1) from days 3 to 6 in generation 1 at sodium concentrations of 60, 200, 250, and 300 mM. **F)** Fecundity at day 6 in generation 1 of the five *C. elegans* strains at each sodium concentration. **G)** Fecundity at day 6 in generation 2 of the five *C. elegans* strains at each concentration. Data is represented as means. Statistical analysis was performed using two-way ANOVA followed by post-hoc Tukey’s HSD tests in PRISM.

In the second generation, all strains showed similar changes in fecundity in response to salt, with significant decreases in fecundity between the control and the different NaCl concentrations, but no significant differences between 200, 250 or 300 mM NaCl (Fig. 5G). The observations for the first generation confirm the existence of negative epistasis between K120M and N122H, where each substitution alone improves salt tolerance, but their combination does not. In this assay, the combination of substitutions even decreased nematode salt tolerance as measured by fecundity relative to nematodes in which the substitutions were present individually, which marks an exacerbation of the negative epistasis observed for the survival assay (Fig. 4A).

## 4. Discussion

Herbivorous insects feed on plants that typically offer relatively low sodium levels. To satisfy their salt requirements many of these insects regularly visit the soil, making them more vulnerable to attack by soil-inhabiting predators and parasites such as EPNs (Xiao et al., 2010; Kaspari, 2020). EPNs are influenced by environmental cues—such as salt, CO_J, plant root- or host insect-derived chemicals, and other factors—when attempting to locate these potential prey insects (Achi et al., 2025a; Dillman et al., 2012b; Nielsen et al., 2011; Rasmann et al., 2005; Zhang et al., 2021). Soil salinity levels can thus impact multi-trophic interactions in soil foodwebs. With human activities directly or indirectly driving increases in soil salinity across large geographic areas, understanding mechanisms of acclimation and adaptation to salt in EPNs is important.

Environmental salt levels can vary significantly, from 1 mM to 300 mM, depending on whether the soil is non-saline or saline (Department of Primary Industries and Regional Development, Western Australia). While information is limited, EPNs, as limno-terrestrial animals (Grant and Villani, 2003), are believed to thrive in low-to-moderate salt concentrations of around 80 mM (Nielsen et al., 2011; Stuart et al., 2015). This could explain why the three *Steinernema* EPN species tested here achieved their highest infection rates at soil salinity levels of 50 mM. Under what seem to be close to optimal conditions (50 mM), *S. feltiae* showed the highest infection rates among the EPN species, significantly exceeding those of *S. carpocapsae* at first, but then dropping sharply at higher salt concentrations. While *S. hermaphroditum* displayed salt tolerance in survival assays, its ability to successfully infect host insects and complete its life cycle was greatly diminished at sodium levels exceeding 150 mM. Although 150 mM may be higher than current salinity levels of soils in most areas, these results offer valuable insights into the challenges EPNs may face when being forced to acclimate or adapt to rising soil salinity levels.

The constraints that increase in soil salinity may pose to different EPN species were further borne out in preference assays in agar or soil media. On agar plates, *S. carpocapsae* exhibited a significant preference for 150 mM over 50 mM NaCl, while *S. hermaphroditum* showed no preference, and *S. feltiae* preferred 50 mM, consistent with their respective salt tolerance levels in survival assays. When tested in sand, *S. feltiae* initially preferred 50 mM NaCl over the control but then began to prefer lower over higher concentrations when NaCl levels exceeded 50 mM. Furthermore, *S. hermaphroditum* always seemed to prefer the control over test options offering different sodium levels. Finally, although *S. carpocapsae* preferred the control over both the 50 mM and 150 mM test options, this species demonstrated significantly higher navigation towards the 300 mM test option compared to the control. These findings suggest that, compared to its congeners, *S. carpocapsae* appears to prefer soils with relatively high salt concentrations, which may be adaptive if it has a competitive fitness advantage over these other species when locating and infecting host insects in highly saline soils.

The results from the foraging behavior and host infection assays suggested that *S. carpocapsae*’s preferences might indeed be adaptive. Previous studies showed that *S. riobrave* experiences reduced foraging and decreased movement under salt stress (Nielsen et al., 2011). In our assays, although none of three *Steinernema* species we tested exhibited negative effects in host-seeking behavior under salt stress, only *S. carpocapsae* continued to successfully infect waxworm hosts when sodium levels increased. IJs of *S. hermaphroditum* showed a significant decrease in numbers of nematodes entering the host at 300 mM NaCl and IJs of *S. feltiae* showed diminished infectivity starting at 150 mM. These results suggest that although *S. hermaphroditum* and *S. feltiae* can still locate hosts in saline environments, they appear to struggle with successfully infecting them when salt stressed. In addition, *S. carpocapsae* may be seeking out environments where it experiences adaptive competitive advantages over other EPN species, but more work will be necessary to confirm this hypothesis.

Several mechanisms likely influence EPN preferences and performance when experiencing saline environments. *S. carpocapsae* evolved unique amino acid substitutions in the first extracellular loop of the NKA relative to other *Steinernema* EPN species. We previously identified that several plant- and blood-feeding insects have evolved substitutions in the first extracellular loop that may confer heightened tolerance to environmental salt levels (Achi et al., 2025b). For example, *Aedes* spp. mosquitoes evolved Q111L and the species *Ae. albopictus* and *Ae. aegypti* exhibit greater tolerance to elevated salinity levels than many *Anopheles* spp. mosquitoes (Kengne et al., 2019; Bradley, 1987). Furthermore, grasshoppers evolved A119S in combination with Q111L and are known to be capable of living on plants in salt-affected areas such as salt marshes (Haines and Montague, 1979; Li et al., 2023). Using functional genetic assays with gene-edited *Drosophila* CRISPR knock-in mutants we found that these substitutions could confer increased salt tolerance, allowing flies to pupate and reach adulthood more successfully in highly saline conditions (Achi et al., 2025b).

We took a similar approach to identify if substitutions K120M and N122H that evolved in *S. carpocapsae* specifically might explain differences in salt tolerance between *Steinernema* species by knocking in these substitutions in *C. elegans*, whose sequence of the first extracellular loop of the NKA is identical to that of *S. feltiae* and *S. hermaphroditum*. In one assay, we exposed nematodes to different agar media containing excess Na□ or K□ to assess their abilities to survive (Khanna et al., 1997). Our results for the N2 wild-type *C. elegans* strain in this assay were similar to those from a previous study (Lamitina et al., 2004), which validates our findings for this strain and our knock-in mutant strains. However, we also performed a second assay in which nematodes were immersed in media with different sodium concentrations and both survival and fecundity fitness were tracked across two generations. This second assay may offer a more reliable assessment of survival as the environment mimicked environments that these limno-terrestrial animals occur in more naturally (Grant and Villani, 2003). From these assays a picture emerged in which substitutions K120M and N122H individually conferred higher salt tolerance to *C. elegans*. Surprisingly, when the substitutions were present in *C. elegans* together, as in *S. carpocapsae*, this advantage disappeared. These findings suggest that the combination of substitutions K120M and N122H in *S. carpocapsae* do not explain this species’ enhanced salt tolerance relative to the other *Steinernema* species and that other factors are likely contributing as well. This latter notion is further supported by our finding that both *S. carpocapsae* and *S. hermaphroditum* exhibited salt tolerance in terms of maintaining survival in the first assay. Future investigations may find further evidence for this conclusion once gene editing becomes possible in EPN model systems.

The pattern of negative epistasis between substitutions K120M and N122H also suggests that although ancestors of *S. carpocapsae* may have experienced improved salt tolerance when the first substitution evolved—this may be either of the substitutions since both evolved along the lineage leading up to *S. carpocapsae*—in the current situation where this EPN possesses both substitutions it is unlikely that sodium levels in the soil represent the agent of selection that may have driven the evolution of the combination of substitutions. We previously identified that *S. carpocapsae* is more infectious than *S. feltiae* and *S. hermaphroditum* on insect herbivores that sequester toxic CGs from milkweeds (Achi et al., 2025a). Although it is currently unknown if substitutions K120M and N122H confer resistance to CGs in nematodes, N122H is known to have a large effect on mediating CG resistance through target-site insensitivity in the monarch butterfly and other insect herbivores that specialize on milkweeds (Karageorgi et al., 2019). Interestingly, the monarch appears to perform well on milkweed plants growing in roadside environments, where milkweed plants may receive relatively high levels of sodium from road salt application—resulting in increased muscle mass in males and enhanced neural investment in females (Snell-Rood et al., 2014). Santiago-Rosario et al. (2024) further found that both male and female monarchs accumulated high sodium levels under elevated Na^+^ treatments, whereas other butterfly species showed significantly higher sodium accumulation only in males. However, when testing *Drosophila* knock-in mutants with the combination of substitutions in the NKA’s first extracellular loop found in the monarch, which includes N122H, we identified no increase in salt tolerance relative to control flies (Achi et al, 2025b). These results suggest an interesting parallel between insects and nematodes: while certain substitutions in the first extracellular loop of the NKA may elevate salt tolerance, evolution of combinations of substitutions involving N122H may instead be driven more by selection mediated by CGs. More research will be needed to evaluate this possibility.

In summary, in this study we identified: (1) that *S. carpocapsae* displays a preference for environments with higher salt concentrations than *S. feltiae* and *S. hermaphroditum*; (2) that *S. carpocapsae* maintains higher levels of infectivity than the other two species in highly saline environments; (3) that substitutions K120M and N122H found in *S. carpocapsae* do not explain differences in salt tolerance between the species, despite a positive effect of each individual substitution on salt tolerance; and (4) that soil salinity levels are unlikely to have driven the evolution of the two substitutions in *S. carpocapsae* since negative epistasis was apparent between them. Further investigation into the mechanisms that underpin salt tolerance in *S. carpocapsae* could deepen our understanding of how this trait impacts multi-trophic interactions in different environmental settings and may reveal how salt tolerance evolved in this species, which could have important ramifications for biological control of insect pests.

## Supporting information

Raw Data

## Acknowledgements

The authors thank Naomi Pierce (Harvard University) for helpful discussions and Larry Wayne Duncan (University of Florida) for help with designing the olfactometers that were used. This study was supported by the National Institute of General Medical Sciences of the National Institutes of Health (award no. R35GM151194 to SCG and award no. R35GM137934 to ARD), the United States Department of Agriculture’s National Institute of Food and Agriculture (award no. 1032189 to ARD), the United States Department of Agriculture’s National Institute of Food and Agriculture Multistate Project (Project W4185, of which ARD is a member), and startup funds from the University of California Riverside (to SCG).

---

**Supplementary Fig. 1:**
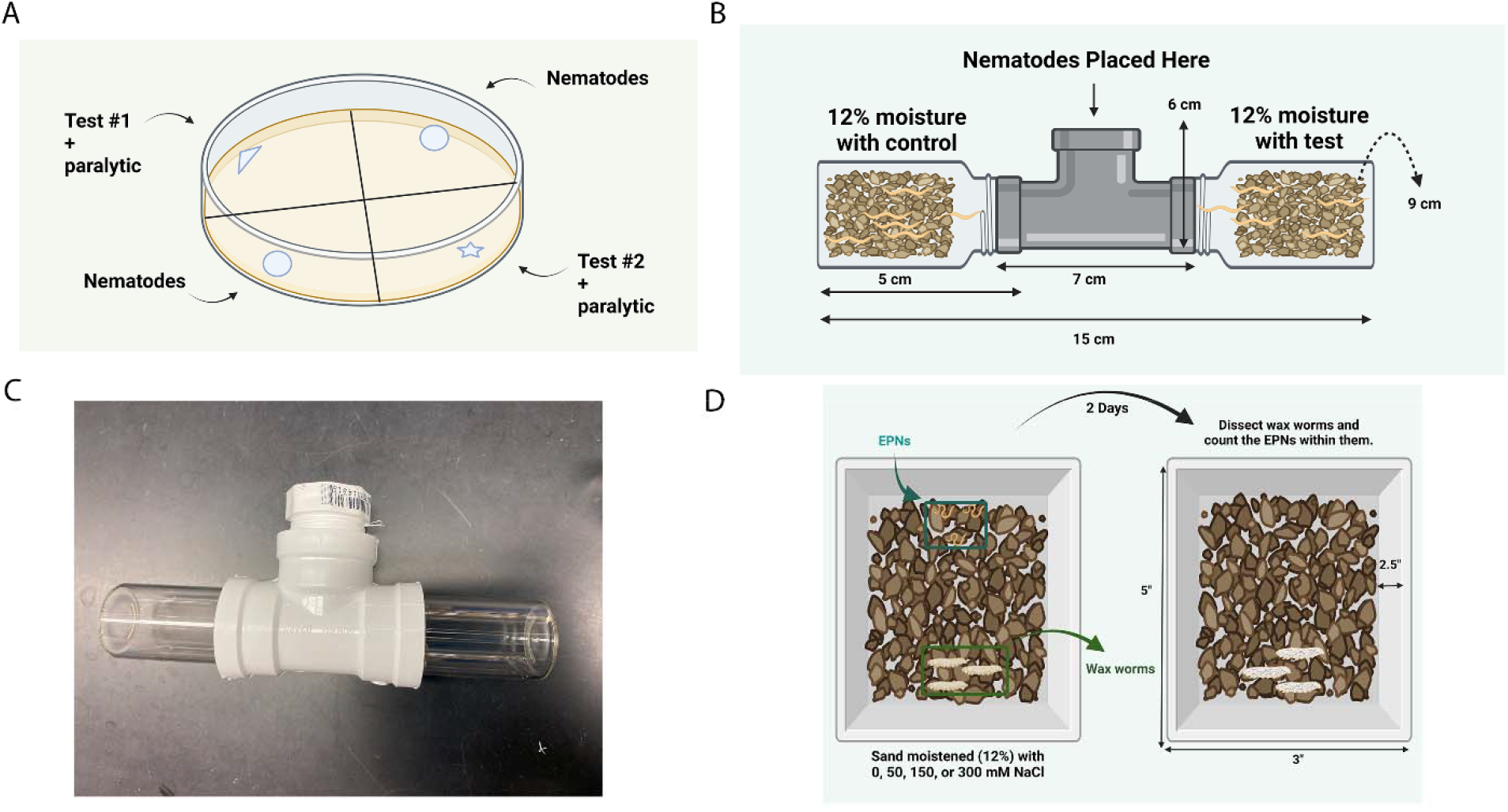
Set ups for three chemotaxis and infectivity assays with entomopathogenic nematodes on agar and in sand. **A)** Chemotaxis assay on agar to test nematode preference for different salt concentrations. **B)** Chemotaxis assay to test the same preference in sand. **C)** Olfactometer for the chemotaxis assay in sand. **D)** Infectivity assay in sand.

**Supplementary Fig. 2:**
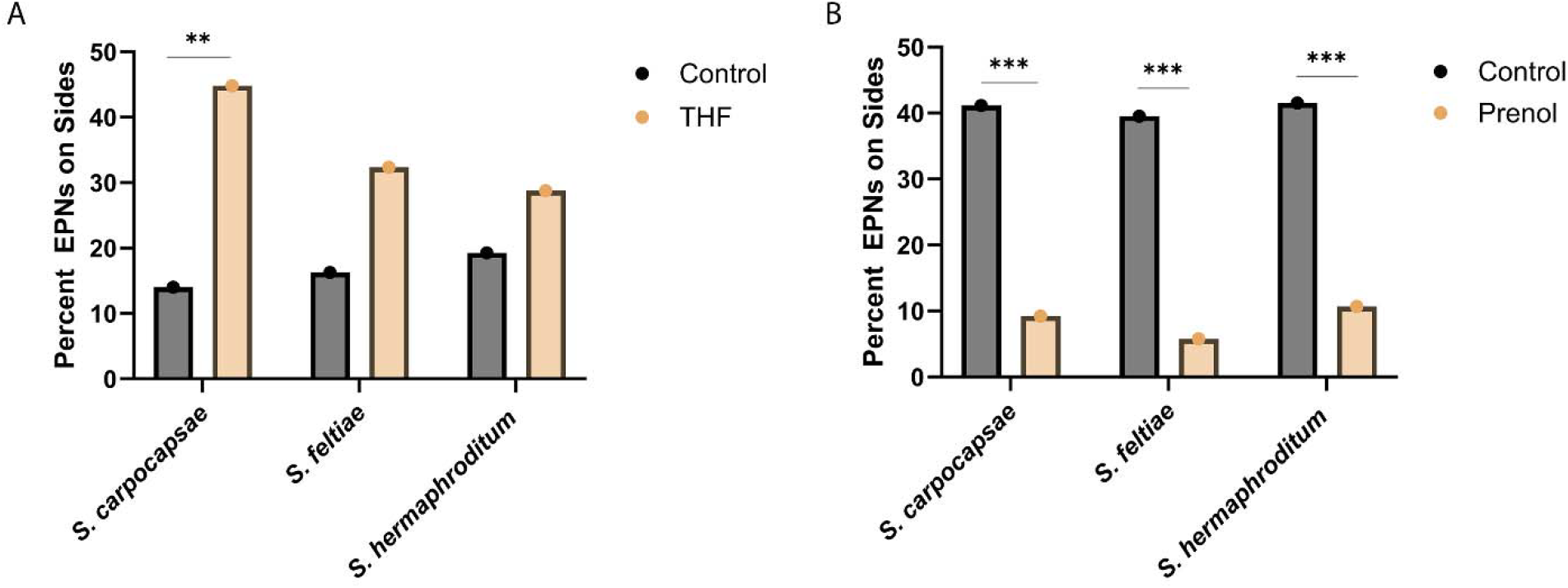
Entomopathogenic nematodes participated in chemotaxis assays on agar as expected based on their behavior towards known attractants and repellants. **A-B)** Percentage of *Steinernema carpocapsae, S. feltiae*, and *S. hermaphroditum* nematodes (respectively) choosing control and test sides in a preference assay with different test solutions: Control vs. THF and Control vs. Prenol. Data is represented as means. Statistical analysis was performed using two-way ANOVA followed by post-hoc Tukey’s HSD tests in PRISM.

**Supplementary Fig. 3:**
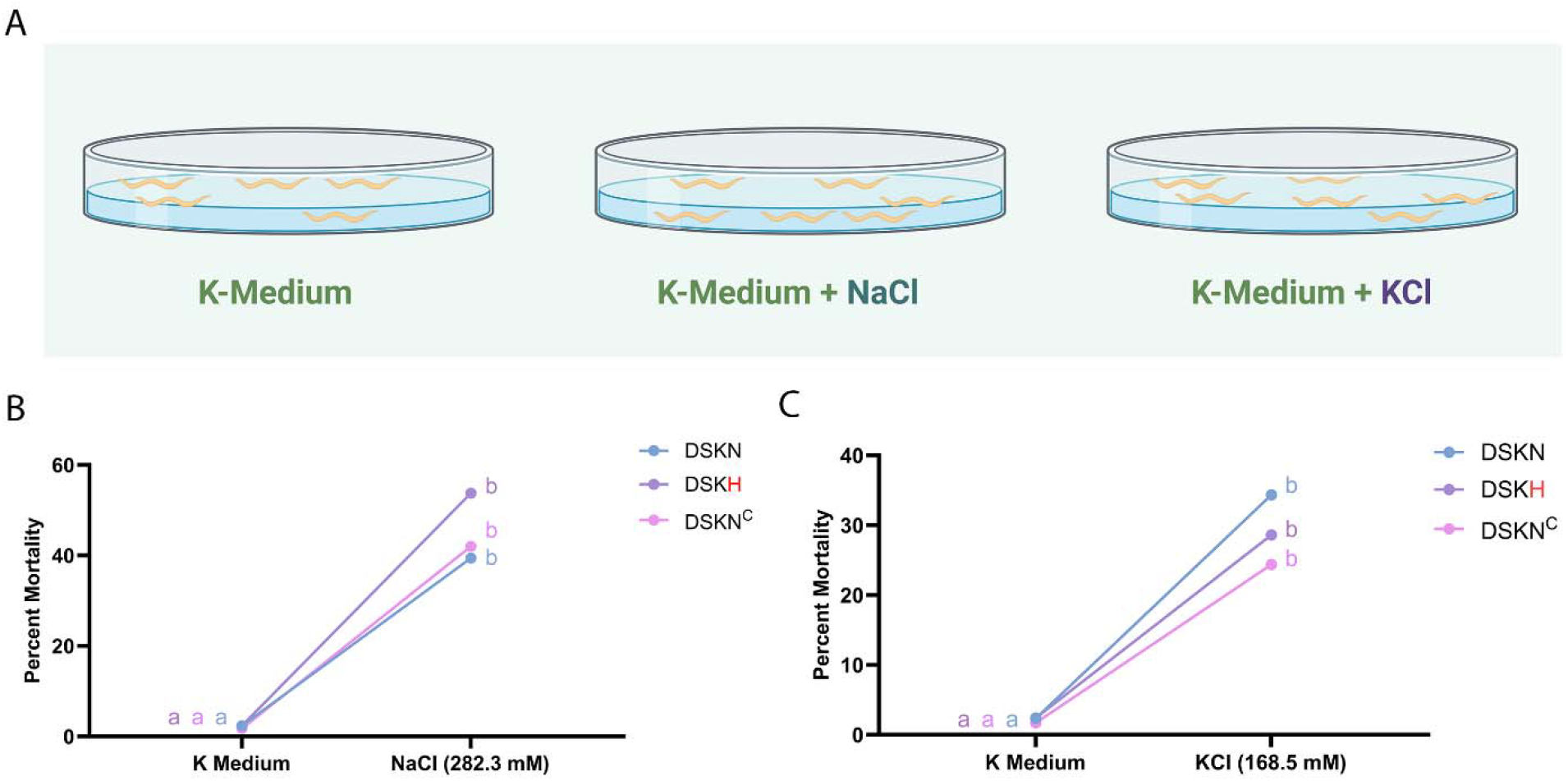
*Caenorhabditis elegans* strains DSKN, DSKN^C^, and DSKH show sensitivity to sodium at relatively high concentrations. **A)** Set up of the salt tolerance assay with the *C. elegans* strains (described in Table 1) and different *Steinernema* species. **B-C)** Percent mortality of *C. elegans* strains DSKN, DSKN^C^, and DSKH on K-medium buffer with and without additional NaCl or KCl, respectively. Data is represented as means. Statistical analysis was performed using two-way ANOVA followed by post-hoc Tukey’s HSD tests in PRISM.

**Supplementary Fig. 4:**
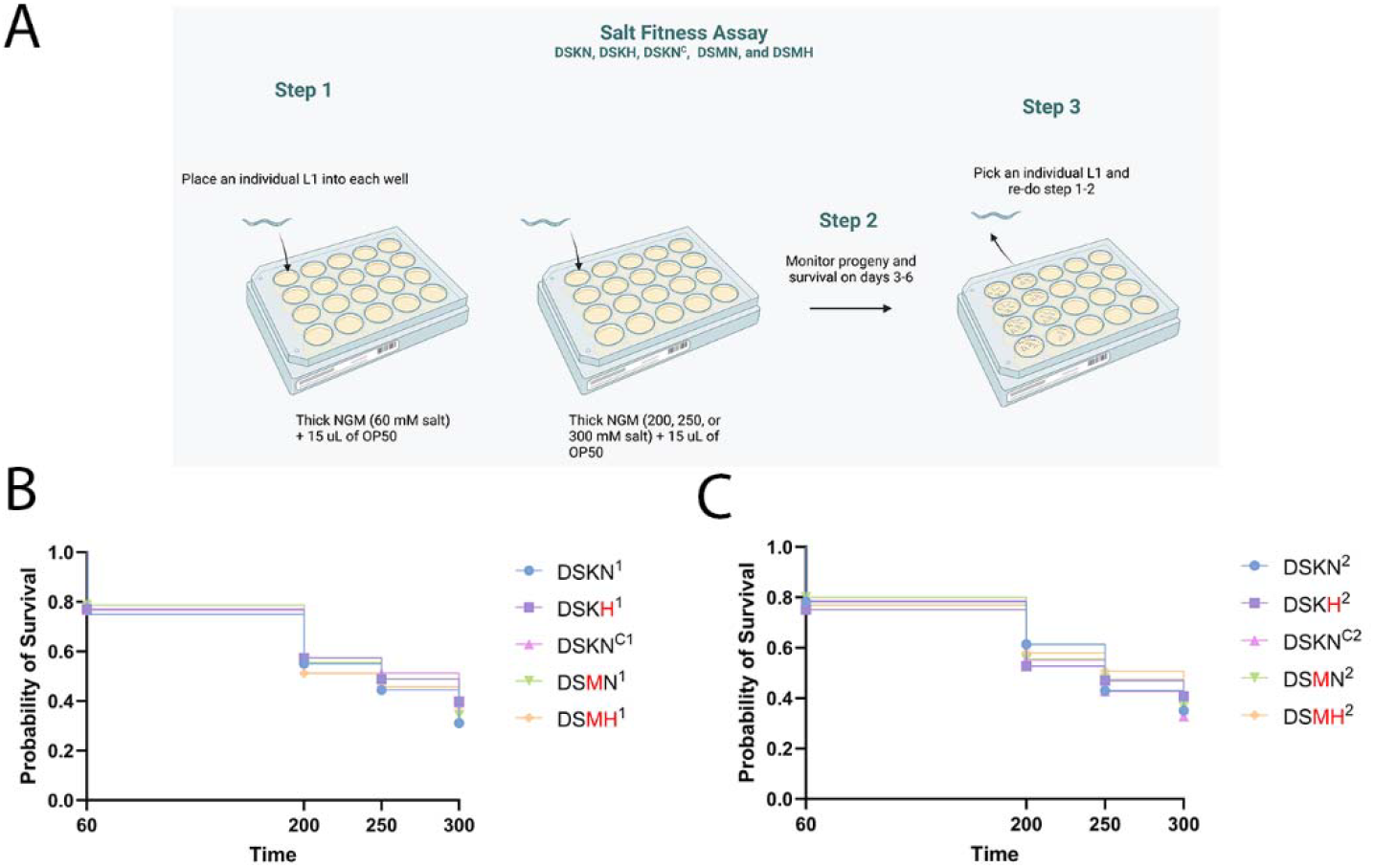
Experimental design of a fitness assay presenting nematodes with different salt concentrations and analysis of survival probabilities. **A)** Set up of salt fitness assay monitoring survival and fecundity of *Caenorhabditis elegans* strains. **B-C)** Probability of survival for generation 1 and generation 2 (consecutively) of *C. elegans* strains at salt concentrations of 60, 200, 250, and 300 mM. Statistical analysis was performed using log-rank tests for survival.

